## Supplemental Tables and Figures for "Transcriptomics reveals the putative mycoparasitic strategy of the mushroom *Entoloma abortivum* on species of the mushroom *Armillaria*" (from another version of this preprint: version 1, 2021-05-01)

**Table S1** Location and GenBank accession numbers of *Armillaria mellea* sequences used in the phylogenetic analysis

| Country | GenBank | Collection/Herbarium no. | Reference |
|---|---|---|---|
| Azerbaijan | EF637086 | B1458 | 1* |
| China | KF032530 | DF1 | 2* |
| China | KF032532 | SY1 | 2* |
| China | KF032534 | DJ4 | 2* |
| China | KF032535 | KY5 | 2* |
| China | KF032536 | KY4 | 2* |
| China | KJ643339 | HZ4 | 2* |
| China | KP162319 | Lijiang | 3* |
| China: Ghizhou | KT822253 | HKAS86592_G_02001_5 | (86) |
| China: Hubei | KT822252 | HKAS86591_G_01010_1 | (86) |
| China: Sichuan | KT822245 | HKAS49004 | (86) |
| China: Yunnan | KT822242 | HKAS85471 | (86) |
| China: Yunnan | KT822243 | HKAS86612_K_00109_4 | (86) |
| China: Yunnan | KT822249 | HKAS85599 | (86) |
| China: Yunnan | KT822251 | HKAS86590_G_00020_6 | (86) |
| China: Yunnan | KT822255 | HKAS86597_G_99044_2 | (86) |
| China: Yunnan | MG931746 | CBSK_CFCC81073 | 4* |
| China: Yunnan | MG931782 | HMAS253414 | 4* |
| China: Zhejiang | KT822244 | HKAS86594_G_04055_9 | (86) |
| China: Zhejiang | KT822254 | HKAS86593_G_04054_14 | (86) |
| Denmark | MG931762 | H7000666 | 4* |
| France | AF163584 | B527 | (87) |
| France | AF163585 | B525 | (88) |
| France | AF163586 | B1245 | (87) |
| France | HQ232290 | D4 | 5* |
| France | JN657464 | D1 | 5* |
| France | JN657465 | D5 | 5* |
| France | MG931767 | H7031840 | 4* |
| France | MN660639 | SREF2371 | (89) |
| Hungary | AF163581 | B1212 | (88) |
| Iran | AF163583 | B1205 | (88) |
| Italy | FJ716618 | CRA-PAV2824 | 6* |
| Italy | FJ875692 | CMW 11265 | (90) |
| Italy | JF907776 | 8602 | (91) |
| Italy | KP960536 | ER1960 | 7* |
| Italy | KP960537 | ER1961 | 7* |
| Italy | KP960538 | ER1962 | 7* |
| Italy | KP960539 | ER1963 | 7* |
| Italy | KP960550 | ER1981 | 7* |
| Italy | KP960552 | ER1983 | 7* |
| Italy | KP960554 | ER1985 | 7* |
| Italy | KP960555 | ER1986 | 7* |
| Italy | KP960556 | ER1987 | 7* |
| Italy | KP960557 | ER1988 | 7* |
| Italy | KP960558 | ER1989 | 7* |
| Italy | KP960561 | ER1992 | 7* |

**Table S1** *continued*

| | | | |
|---|---|---|---|
| Italy | KP960562 | ER1993 | 7* |
| Italy | KP960563 | ER1994 | 7* |
| Italy | KP960564 | PF1551 | 7* |
| Italy | KP960565 | ER1996 | 7* |
| Italy | KP960566 | ER1997 | 7* |
| Japan | AF163594 | B731 | (88) |
| Japan | KT822246 | HKAS86588_83003_2 | (86) |
| Japan | MF095794 | TNS:F-70421 | 8* |
| Poland | AY848938 | UASWS0027 | 9* |
| Poland | KX756392 | WA0000052261 | (92) |
| Portugal | KP960535 | ER1959 | 7* |
| Scotland | JF313749 | TFB4184 | (93) |
| Slovenia | AJ250051 | M1 (90260/1) | (94) |
| Slovenia | KF020706 | 17_7_9 | 10* |
| South Korea | AF163591 | B608 | (88) |
| South Korea | AF163592 | B916 | (88) |
| South Korea | AF163593 | B917 | (88) |
| Spain | EU266545 | AH9328 | (95) |
| Spain | EU266546 | GDA51999 | (95) |
| Sweden | KY352514 | olrim1073 | 11* |
| Tibet | MG931786 | HMAS253585 | 4* |
| Tunisia | KU863550 | Oe8 | 12* |
| United Kingdom | AF163578 | B176 | (88) |
| United Kingdom | AF163579 | B1247 | (87) |
| United Kingdom | AF163580 | B1240 | (87) |
| United Kingdom | AF163582 | B186 | (87) |
| United Kingdom | KX618568 | MCOLLINS12 | 13* |
| United Kingdom | KX618569 | MCOLLINS15 | 13* |
| United Kingdom | KX618581 | MK7 | 13* |
| United Kingdom | MH855148 | CBS 106.31 | (96) |
| Ukraine | JN657466 | HY3 | 5* |
| USA | FJ664593 | VI 49.5.1 | 14* |
| USA: California | AF163595 | B927 | (88) |
| USA: California | AF163596 | B931 | (88) |
| USA: California | AF163597 | B1217 | (88) |
| USA: California | FJ664595 | PVI 85.43.1 | 14* |
| USA: California | FJ664596 | PVI 84-45.1 | 14* |
| USA: Georgia | MG931920 | TENN054336 | 4* |
| USA: Massachusetts | AF163589 | B496 | (88) |
| USA: Massachusetts | AF163590 | B497 | (88) |
| USA: Massachusetts | AY789081 | PBM2470 | (97) |
| USA: Massachusetts | MF161188 | FH:BHI-F169 | (98) |
| USA: Minnesota | MG931908 | MIN0877204 | 4* |
| USA: New Hampshire | AF163587 | B282 | (88) |
| USA: New Hampshire | AF163588 | B623 | (88) |
| USA: New Hampshire | AY213587 | ST21 | (99) |
| USA: New Hampshire | KT822256 | HKAS86611_B275 | (86) |
| USA: South Carolina | AY526602 | GA.Apple clone 1 | (100) |
| USA: South Carolina | AY526603 | SC.JC-3.02 clone 1 | (100) |
| USA: South Carolina | AY526604 | SC.JC-3.02 clone 2 | (100) |

**Table S1** *continued*

| | | | |
|---|---|---|---|
| USA: South Carolina | MG931929 | TENN061702 | 4* |
| USA: Tennessee | MF755264 | JMC038 (TENN) | 15* |
| USA: Virginia | AY213584 | ST5-A | (99) |
| USA: Virginia | AY213585 | ST5-B | (99) |
| USA: Wisconsin | AY213586 | ST20 | (99) |
| Outgroup | | | |
| *Armillaria gemina* | FJ664586 | II JB-38A.1 | 14* |
| *Armillaria sinapina* | FJ664609 | V 48.5.1 | 14* |
| *Armillaria puiggarii* | KU170954 | MCA 3111 | (13) |
| *Armillaria luteobubalina* | FJ664591 | V M 5.2.1 | 14* |

1*: Aghayeva DN, Harrington TC, unpublished
2*: Huang W, Zhu G, Liu C, unpublished
3*: Guo S-X, unpublished
4*: Liang J, Pecoraro L, and Cai L, unpublished
5*: Tsykun T, Prospero S, Rigling D, unpublished
6*: Scire, Motta and D'Amico, unpublished
7*: Tizzani L, Haegi A, Motta E, unpublished
8*: Kasuya,T., Maruyama,T., Fuse,K., Hosaka,K. and Minowa,K., unpublished
9*: Calmin,G., Belbahri,L. and Lefort,F., unpublished
10*: Piskur B, Hauptman T, unpublished
11*: Smith,G.R., Finlay,R., Stenlid,J., Vasaitis,R. and Menkis,A., unpublished
12*: Gharbi Y, unpublished
13*: Denman,S., Barett,G., Kirk,S.A., McDonald,J.E. and Coetzee,M.P.A., unpublished
14*: Stefani FOP, Berube JA, Hamelin RC, unpublished
15*: Matheny,P.B. and Birkebak,J.M., unpublished
