## Supplementary material for "Transcriptomics reveals the putative mycoparasitic strategy of the mushroom *Entoloma abortivum* on species of the mushroom *Armillaria*": Cover Letter for mSystems submission (from another version of this preprint: version 1, 2021-05-01)

**RACHEL A. KOCH, PH.D.**
Postdoctoral Researcher
University of Nebraska
Department of Plant Pathology
Lincoln, NE 68503

30 April 2021

Dear mSystems,

Enclosed, please find the submission file for our manuscript:
**Transcriptomics reveals the mycoparasitic strategy of the mushroom *Entoloma abortivum* on species of the mushroom *Armillaria*.** Herein, we used transcriptomics of field-collected mushroom structures to determine that the mushroom-forming species, *E. abortivum*, is a mycoparasite on mushrooms of *Armillaria*. While mycoparasitism is well-studied in the Ascomycota, this is the first study to analyze the genetics of mycoparasitism in the Basidiomycota. Interestingly, we found that even distantly-related mycoparasites utilize similar genetic mechanisms. Additionally, our understanding of how fungal species interact with one another in nature is nascent, and this work represents a large step forward in that understanding. Because of the microbial system that is the focus of this paper, and the methodologies we used to derive our conclusions, we believe this paper is well-suited for publication in this journal.

Appropriate reviewers for this manuscript include: Dr. Alisha Quandt (University of Colorado, Boulder;), Dr. Daniel Lindner (US Forest Service;) and Dr. Magnus Karlsson (Swedish University of Agricultural Sciences;).

If this manuscript does go out for review, we will be sure to have accession numbers for the transcriptomes and raw read data in time for revisions. We thank you for your consideration and look forward to hearing from you! Please let us know if there is anything else we need to provide in the meantime.

Sincerely,

Rachel A. Koch, Ph.D.

425 Plant Sciences Hall ■ 1875 N 38th St. ■ Lincoln, NE 68503 ■ (320) 260-3702
